# Are Sources of EEG and MEG rhythmic activity the same? An analysis based on BC-VARETA

**DOI:** 10.1101/748996

**Authors:** Usama Riaz, Fuleah A. Razzaq, Deirel Paz-Linares, Ariosky Areces-Gonzalez, Sunpei Huang, Eduardo Gonzalez-Moreira, Maria L. Bringas Vega, Eduardo Martinez Montes, José Enrique Alvarez Iglesias, Pedro A. Valdés-Sosa

## Abstract

In the resting state (closed or open eyes) the electroencephalogram (EEG) and the magnetoencephalogram (MEG) exhibit rhythmic brain activity is typically the 10 Hz alpha rhythm. It has a topographic frequency spectral distribution that is, quite similar for both modalities--something not surprising since both EEG and MEG are generated by the same basic oscillations in thalamocortical circuitry. However, different physical aspects underpin the two types of signals. Does this difference lead to a different distribution of reconstructed sources for EEG and MEG rhythms? This question is important for the transferal of results from one modality to the other but has surprisingly received scant attention till now. We address this issue by comparing eyes open EEG source spectra recorded from 70 subjects from the Cuban Human Brain Mapping project with the MEG of 70 subjects from the Human Connectome Project. Source spectra for each voxel and frequencies between 0-50Hz with 100 frequency points were obtained via a novel sparse-covariance inverse method (BC-VARETA) based on individualized BEM head models with subject-specific regularization parameters (noise to signal ratio). We performed a univariate permutation-based rank test among sources of both modalities and found out no differences. To carry out an unbiased comparison we computed sources from eLORETA and LCMV, performed the same permutation-based comparison, and found the same results we got with BC-VARETA.

## 1. Introduction

At resting state EEG and MEG readings show an evident rhythmic activity in frequency spectra, specifically alpha band. Postsynaptic potentials (PSP) are continuously happening even in resting state (both eyes open and closed). These PSPs are generated from the same cortical networks (thalamocortical, cortical-cortical, etc.) PSPs generate primary current densities (PCD) at the cortical surface and these PCDs are measured as electric potential or magnetic field by EEG electrodes and MEG magnetometers respectively. Since both phenomena are generated from the same cortical activity, there should not be any differences in these rhythms which seems evident at first sight. However, both modalities are physically different, so they suffer from different effects of volume conduction. Moreover, the measurement noise in the recorded signals might be different since the engineering and physics behind the signal and sensors that are used for acquiring those signals are different for both modalities.

These biological and instrumental factors affect the M/EEG signals. However, it’s an ongoing discussion that whether the sources of the acquired rhythmic activities are the same or different. The term source localization is used to define the process of reconstructing the cortical source spectra or to localize the cortical activity from the electromagnetic rhythms. While comparing source localization, there are different results based on a variety of experiments. Some claimed EEG source localization is better than MEG (Liu, Dale, & Belliveau, 2002), (Gavaret, Badier, Bartolomei, Bénar, & Chauvel, 2014) (Klamer, et al., 2014) others found in their experiments that MEG has better source localization accuracy (Cohen & Cuffin, 1991) (Cuffin, 1983), while others did not find significant differences (Hedrich, Pellegrino, Kobayashi, Lina, & Grova, 2017) (Waldert, et al., 2008) (Cuffin, 1983) between two modalities performance on source localization. There is a significant amount of research that claims combining EEG and MEG outperforms individual modality in terms of source localization and spike detection (Lin, et al., 2003) (Knake, et al., 2006) (Sharon, Hämäläinen, Tootell, Halgren, & Belliveau, 2007) (Muthuraman, et al., 2014) (Plummer, et al., 2019).

One major difference between EEG and MEG is the sensitivity to source orientation and source Signal-to-Noise Ratios (SNR) at different brain areas. Evidence for the sensitivity of EEG to tangential and MEG to radial sources are found in many studies (Cuffin, 1983) (Haueisen, Funke, Güllmar, & Eichardt, 2012). However, there are results showing the opposite case and that opens the discussion of the sensitivity of EEG and MEG to different source orientation (Hunold, Funke, Eichardt, Stenroos, & Haueisen, 2016) (Rossi, Luria, Sommariva, & Sorrentino, 2017). The sensitivity of EEG and MEG to deep and superficial sources has been discussed in many studies claiming MEG is not able to give high SNRs for deep sources while EEG is successful in that (Hunold, Funke, Eichardt, Stenroos, & Haueisen, 2016).

As mentioned earlier that volume conduction effect has an influence on the readings of both modalities. This is demonstrated in the experiments conducted by many researchers for different head models (Vorwerk, et al.) (Siems, Pape, Hipp, & Siegel, 2016). They also found that tissue anisotropy and the white matter has major conduction effect for EEG while MEG is only affected by white matter anisotropy (Haueisen, et al., 2002) (Siems, Pape, Hipp, & Siegel, 2016). These two phenomena also affect source reconstruction accuracy where EEG is more susceptible to muscle artifacts and different head models while MEG is less prone to both factors (Wolters, et al., 2006).

All of the comparative studies to date do not compare EEG and MEG statistical analysis of source spectra, which is the main highlight of this study Additionally, there are some notable shortcomings in almost all the current studies. For example; real head models were not used in most studies, the number of subjects was not significantly large, modern inverse solutions were not used to incorporate the effects of cortical activity as well as connectivity estimation techniques. To carry out a comprehensive and conclusive comparative study between two modalities, it is very important to consider all the above define aspects. This study has been carried out by comparing the resting state (eyes open) of 70 subjects from Cuban Human Brain Mapping project and MEG of 70 Human Connectome Project. Figure 1. shows a flowchart for the methodology that has been used for statistical comparison of two modalities. We have used a novel inverse method BC-VARETA and compare the results achieved with other inverse methods e.g. eLORETA and LCMV. We perform statistical testing on the results of the inverse solutions by carrying out multiple comparisons using permutations test and found both modalities don’t have any differences. These results open a whole new field of conducting studies that were limited to only either of the modalities and also to carry out cross-modality source spectra transferal.

**Figure 1:**
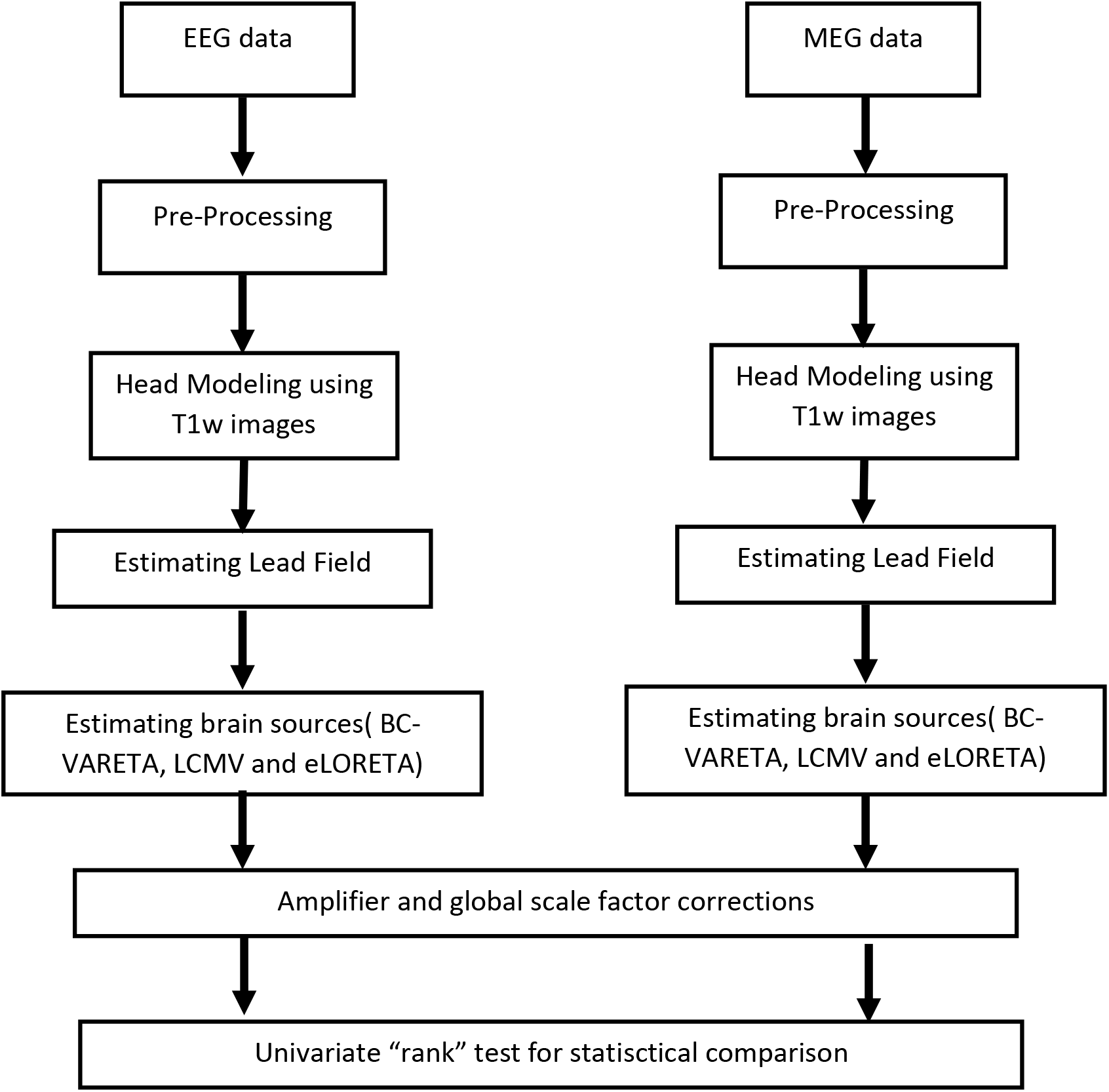
Flow chart for methodology

## 2. Materials and Methods

### 2.1. Dataset

To perform the analysis, an equal number of subjects were used from two big databases i.e. Cuban Human Brain Mapping (CHBM) project and Human Connectome Project (HCP). The EEG data from the Cuban Brain Mapping Project (Hernandez-Gonzalez, et al., 2011) was recorded using 58 channels, electrodes were placed according to the international 10/20 electrode system. This data was acquired from 70 healthy subjects (60 male and 10 females, ages between 19 to 50 years old) in resting state with eyes open. The EEG amplifiers used while gathering this data have a transfer function similar to a bandpass filter of 0-50Hz. So, the EEG data gathered using these amolifiers was filtered at 0 - 50 Hz. In the end, Independent Component Analysis (ICA) (Comon, 1994) was applied to the time-series data to remove the artifacts. The sampling frequency was 200 Hz. Magnetic resonance imaging (MRI) was performed on (MAGNETOM Symphony Siemens, 1.5 Tesla) equipped with a 32-channel head coil.

The MEG data that has been studied in this work is from the Human Connectome Project (Van Essen, et al., 2011) (Behrens, et al., 2013)(Van Essen et al., 2011) (Behrens et al., 2013), led by Washington University, University of Minnesota, and Oxford University (https://www.humanconnectome.org/) (Hileman, et al., 2013). The MEG data were acquired using 248 magnetometer channels and 23 reference channels of a whole head MAGNES 3600 (4D Neuroimaging, San Diego, CA) system. The sampling frequency of the preprocessed data was 508 Hz. We analyzed 70 healthy subjects (37 males and 33 females, ages between 22 to 35 years old) with eyes open restingstate condition. The MEG data were preprocessed by the Human Connectome Project teams using MEG Connectome pipeline (Van Essen, et al., 2011). MEG amplifiers have a transfer function equivalent to a bandpass filter of 1.3-150 Hz and two additional notch filters of 59-61 Hz and 119-121 Hz. Moreover, ICA was used for artifact removal. MRI for the anatomical data was collected using a 3 Tesla (3T) Siemens Skyra scanner.

### 2.2. Forward model

To estimate the realistic head models and estimation of the lead field generation for each subject, a pipeline based on the Freesurfer toolbox, Ciftify (HCP) (http://www.freesurfer.net/) (Dale, Fischl, & Sereno, 1999) (Fischl, 2012) (Dale, Fischl, & Sereno, 1999) (Fischl, 2012) and Brainstorm Toolbox (https://neuroimage.usc.edu/brainstorm/) (Tadel, Baillet, Mosher, Pantazis, & Leahy, 2011) (Tadel, Baillet, Mosher, Pantazis, & Leahy, 2011) was used (Fuchs, Kastner, Wagner, Hawes, & Ebersole, 2002). This pipeline is based on three major steps to generate a lead field from T1 raw images while performing some segmentation, denoising, normalization, co-registration of different brain and non-brain regions and electrodes, format transformation, and using some predefined surface registration and generation steps. In the first step, raw T1 MRI images were given as input to the Freesurfer toolbox to perform multiple corrections, segmentation, and reconstructions for cortical, sub-cortical, and non-brain tissues. This was done using the “recon_all” function of the Freesurfer toolbox. The output from the Freesurfer toolbox was given as input to the Ciftify toolbox to perform convert Freesurfer directory into the CIFITFY space or HCP format directory. The same “recon_all” function was used to get this standard output. Later different functions from the Brainstorm tool were used to generate the lead field for 8002 sources. In the Brainstorm toolbox first step was to import non-brain and brain tissues separately and coregister them on each other to generate a complete head model. Later, EEG/MEG electrodes/sensors were imported and coregistered with head models automatically and later visual inspection was done to perform any manually corrections needed. In the end, the Boundary Element Method (BEM) implemented by the OpenMEEG toolbox embedded in the Brainstorm software was used to generate Lead Field.

To compute the Lead Field of both EEG and MEG, we used a structural pipeline [10] based on the Human Connectome Project (HCP) structural processing [1,2] and Brainstorm (BS) [3]. The code is freely available in GitHub (https://github.com/CCC-members/BrainStorm_Protocol). Roughly the steps of this pipeline are: 1-Segmentation of the brain using HCP structural pipeline, this provides MRI T1 images registered linearly to the MNI space, and the corresponding Gray Matter volumetric and superficial spaces used to define the MEG/EEG sources. These spaces are in one-to-one correspondence to the FSAverage volumetric and low resolution (64K) superficial canonical spaces, and therefore allow rapid mapping of the database results to FSAverage used to compare the results. The EEG database contained only MRI T1 images; therefore, we used the Ciftify [4] release of HCP pipelines based on T1 only. This pipeline produced the structural priors used by sSSBL that included cortical parcels, curvature, normal directions, and Laplacian. 2-The head tissue was modeled using FSL [5,6,7] and included scalp, inner skull, and outer skull that was used in BS to define the head model for MEG/EEG Lead Fields. These were extracted from the MRI T1 registered linearly to the MNI space. 3-Computation of the MEG/EEG lead field to a low-level programming pipeline based on BS functionalities such as head modeler, MEG overlapping spheres [8], and EEG OpenMEEG [9]. Several functionalities were included this such as processing specific HCP, Ciftify, and FSL outputs, optimization of processes to produce more accurate registration of the MEG/EEG layouts, open parametrization of the head modeler, and geometric quality control of head model and registration. The latter was used to classify the subjects suitable for MEG/EEG comparison and to perform manual corrections. 4-Moduli for registration of all the results to FSaverage structural space (64K canonical or user-defined) and normalization of solutions.

### 2.3. Source analysis

Since source analysis was performed in the frequency domain, so the frequency spectra of EEG and MEG time-series data was generated for this purpose. To get a complete picture of brain activity, source analysis was performed all frequencies between 0.1-50 Hz with a step size of 0.5 making it a total of 100 frequency points. This was performed the same for both EEG and MEG frequency spectra. The reason for performing source analysis between 0-50Hz was that EEG data was recorded between these frequency ranges (due to amplifier filtering effect). So, source analysis for MEG was also kept limited to these frequency ranges solely for this study. Brain Connectivity Variable Resolution Tomographic Analysis (BC-VARETA) (https://github.com/CCC-members/BC-VARETA_Toolbox) was used to perform source localization. BC-VARETA is a toolbox for the study of MEG/EEG spectral activity and connectivity that, unlike other general MEG/EEG source analysis methods, is designed to minimize the distortions of specific statistical quantities by introducing priors directly on them (Gonzalez-Moreira et al. 2017). Conventional methods make use of a cost function to estimate-for instance— activations that assume certain spatial structure (priors) of the source vectors to regularize the ill-conditioning in the estimation. Then, meaningful statistical quantities of source activity like the second-order statistical moment of the Fourier transform (Spectrum) are obtained as postprocessing of the formerly estimated time series of source activity. The flaw consists in that at the level of spatial distortion in the time-series expected for this type of regularization is not the same as that of the second-order moment -or the Fourier transform. Therefore, the only way to control distortions of the spectrum is by employing a cost function whose regularization term has been placed upon the spectrum and source activation. For this purpose, we used the module of BC-VARETA specifically designed to regularize the spectrum, denominated Spectral Structured Sparse Bayesian Learning (sSSBL) (https://github.com/CCC-members/BC-VARETA_Toolbox/blob/master/functions/activation_level/sSSBLpp.m). The prefix Spectral refers to a modification of SSBL to directly obtain the Fourier coefficients of source activity with minimal distortions of the spectrum, via complex-valued group Elastic-Net (Gonzalez-Moreira et al. 2020). Since BC-VARETA is a novel method, we performed source localization of frequency spectra with two other famous inverse solutions i.e. eLORETA, LCMV for the sake of confirmatory, comprehensive, and comparative analysis.

#### 2.3.1. LCMV beamformer

A spatial filtering method for localizing sources of brain electrical activity from surface recordings is described and analyzed. The spatial filters are implemented as a weighted sum of the data recorded at different sites. The weights are chosen to minimize the filter output power subject to a linear constraint. The linear constraint forces the filter to pass brain electrical activity from a specified location, while the power minimization attenuates activity originating at other locations. The estimated output power as a function of location is normalized by the estimated noise power as a function of location to obtain a neural activity index map. Locations of source activity correspond to maxima in the neural activity index map. The method does not require any prior assumptions about the number of active sources of their geometry because it exploits the spatial covariance of the source electrical activity.

LCMV defines a filter matrix A is employed in a linear transformation from the sensor level to the brain space. A filters the source activity (in a given frequency band or time window) at the ith voxel (grid point) with unit gain while suppressing contribution from the other voxels. The filter depends on the data by means of the covariance.

#### 2.3.2. eLORETA

The eLORETA method is a discrete, three-dimensional (3D) distributed, linear, weighted minimum norm inverse solution. It is also important to emphasize that eLORETA has no localization bias even in the presence of structured noise.

It should be emphasized that the localization properties of any linear 3D inverse solution (i.e. tomography) can always be determined by the localization errors to test point sources. If such a tomography has zero localization error to such point sources located anywhere in the brain, then, except for low spatial resolution, the tomography will localize correctly any arbitrary 3D distribution. This is due to the principles of linearity and superposition. These principles do not apply to non-linear inverse solutions, nor do they apply to schemes that are seemingly linear but are not 3D inverse solutions (e.g. one-at-a-time best fitting dipoles).

### 2.4. Amplifier and global factor corrections

EEG and MEG were recorded in different settings and at different timelines, their amplifiers have different specifications and transfer function. MEG amplifiers have much flatter and wider transfer function (as mentioned in section 2.1) in comparison to EEG amplifiers. This amplifier effect should be corrected as it will affect the inverse solution too, so it will be safe to consider a global factor affecting each individual of all frequencies and all voxels. We will look into the factors affecting the inverse solution spectra.

#### 2.4.1. Amplifiers

A possible source for differences can be a different types of amplifiers form EEG and MEG. EEG and MEG have amplifiers that posses different transfer functions and both of them allow different frequencies to pass. EEG has an amplifier that possesses the property of a bandpass filter of 0-50 Hz. MEG on other hand has the property of a bandpass filter of 1.3-150 Hz and two notch filters of 59-61 Hz and 119-121 Hz. To have a fair comparison we need to correct for these amplifier differences. To do that, we performed the following empirical corrections on the source spectra of EEG and MEG.

Let our original (unfiltered) EEG signal be ^*V_0_(ω)*^ for frequency ^*ω*^. If the EEG amplifier response is *h_v_* (*ω*) the filtered EEG response is

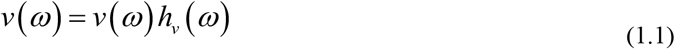

And therefore, the estimated spectrum of the EEG is

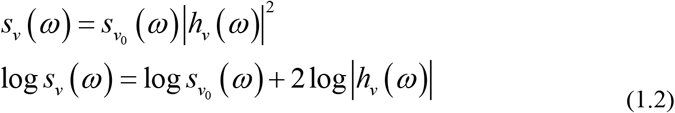

With a similar line of reasoning then one can consider for the MEG

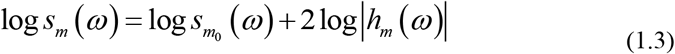

Therefore, the log between EEG and MEG only due to differences in amplifiers is:

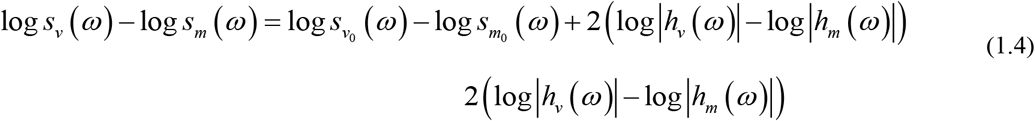

Thus, one can expect a constant difference 2(log*|h_v_* (*ω*)|-log|*h_m_* (*ω*)|) between spectra just due to the different amplifiers. One must adjust for this type of difference between modalities. This scale factor will affect the inverse solutions too So it would be safe to assume that there is a constant amplifier factor that will affect all individuals. For individual ^*i*^, voxel ^*x*^, and frequency ^*ω*^:

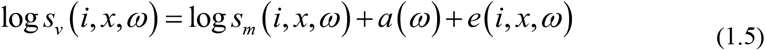

#### 2.4.2. Global-scale factors for EEG (and possibly for MEG)

We have shown previously [1] that EEGs are possibly multiplied by a global scale factor. This would imply an additional, additive, log nuisance parameter that would be constant for all frequencies and voxels over a given individual. Combining this with (1.5) we have:

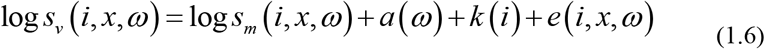

These factors should be corrected before comparing EEG and MEG. It would be extremely easy with a GLM if we had corresponding EEG and MEG for each subject. So, we must look for an ad hoc correction.

#### 2.4.3. Corrective factors

Since the simultaneous EEG and MEG data are not available for each subject, which rules out the possibility of GLM correction also the factors defined in Equation 1.6 are empirically not known. Here, an ad-hoc correction, was adopted to correct for all the above-defined factors. The first correction was made by correcting for any scale differences amongst individuals. This was carried out by defining a factor, *α(i)* that takes mean of log spectra for all voxel(sources) and frequencies. *α(i)* was subtracted from all voxels and all frequencies for within an individual and for all individuals. The second correction was made by defining another factor *β(ω)* for each frequency of EEG/MEG. *β(ω)* was computed by taking the mean for the log spectra of all subjects and all voxels(sources). Factors *α(i)* and *β(ω)* were computed for both EEG and MEG. Once the factors were computed, they were subtracted from the log source spectra of EEG and MEG (to apply the corrections)as shown in Equation 1.7.

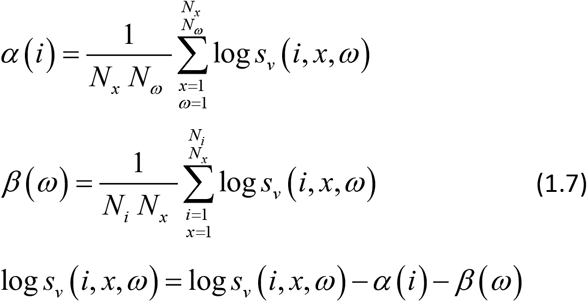

Once the above-defined corrections were made in the log source spectra of EEG and MEG, it can be safely said both frequency spectra have been corrected for any possible differences and they can be compared statistically to look for the underlying physical differences between the modalities.

### 2.5. Statistical Testing for Comparing Sources of M/EEG rhythmic activity

After the necessary corrections were made, the next step was to carry out statistical analysis to find whether two modalities have any or no difference in terms of source spectra. For this purpose we have performed a univariate rank test with multiple permutations using the Flip package in R. We applied this test for all three inverse solutions with 8002 generators/sources and 100 frequency points. Thus, for each subject/individual, we had 800200 variables.

#### 2.5.1. Hyposthesis for zero-inflated data

In case of BC-VARETA, we have divided data into two groups: *G1* is for EEG and *G2* is for MEG. Each group has 800200 variables where each variable is some source s for a specific frequency *f*. When data is highly sparse due to a large number of unobserved values this is known as Zero Inflation. While dealing with source leakage and to minimizes the False Positive sources, BC-VARETA use Sparse Hermitian Sources Graphical Model (Paz-Linares, Gonzalez-Moreira, Martinez-Montes, & Valdes-Sosa, 2018). This generates a large number of zero activations. Thus, the resultant source spectra were highly sparse or Zero-Inflated. The histogram of source activations is shown in Figure 3. which demonstrates zero inflation in reconstructed source space. Thus, the dataset can be modeled as:

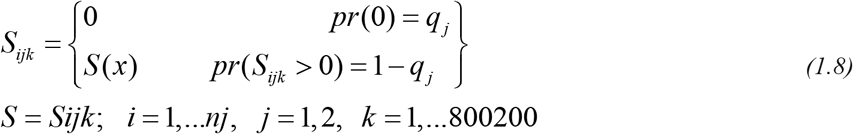

where: *i* is subject, *j* is the sample or group (EEG or MEG), *S(x)* is the observed variable, *S*(*x*) is the underlying distribution of the variables. The univariate null hypothesis to test the similarity between two samples would then be

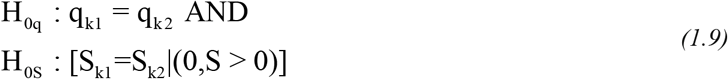

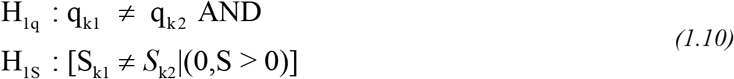

here: ^*H_0q_*^ and ^*H_1q_*^ is the null and alternate hypothesis respectively for the observed zeros where probability ^*q*_1_^ (for sample 1:EEG) should be equal to ^*q*_2_^ (for sample 1:MEG) and k is the response variable. Since BC-VARETA is a sparse inverse solution, there were many vectors with all zero values in both EEG and MEG. To optimize our process, before testing our hypothesis those variables/vectors were removed/omitted before performing any analysis.

**Figure 3:**
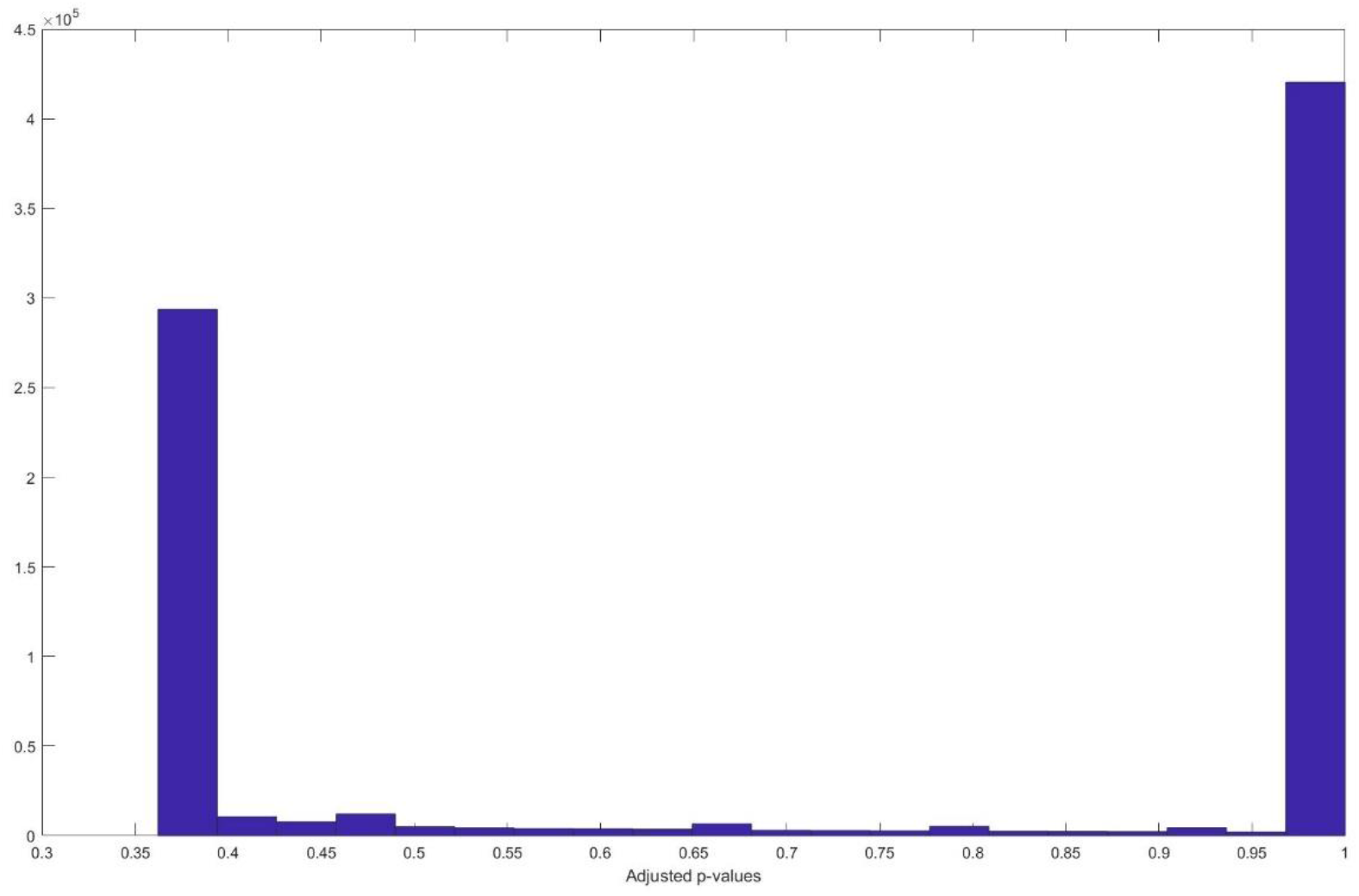
Histogram of adjusted p-values for eLORETA

#### 2.5.2. Hyposthesis for non-zero nflated data

The above defined hypothesis are for the zero-inflated data. However, in case of non-zero inflated source data i.e. eLORETA and LCMV the handling of zero data was not required. Eventually our hypothesis are empirically reduced to:

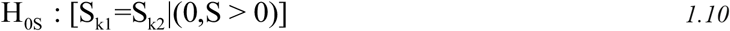

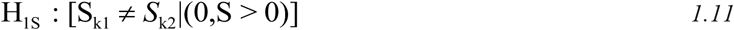

Here, ^*H_0q_*^ and ^*H_1q_*^ no longer exist as they were defining the hypothesis for zeros present in zero-inflated data. Rather, our hypothesis is defined by representing only non-zero data as shown in equations 1.10 and 1.11. Rest of the process for dividing data into groups separately for EEG and MEG and later performing the statistical analysis will remain same.

For above defined scenarios of zero-inflated and non-zero inflated data we have tested the defined hypothesis with univariate permutations based rank test using the FLIP package in R. Permutation or randomization test construct a sampling distribution which in called “permutation distribution” by rearranging and resampling the observed data. In other words, it shuffles or permutes the observed data without replacement and perform some kind of statistical test on each permutation. In this study, we specifically perform the rank test on our data, since our data is not normally distributed. The rank test in the FLIP package specifically uses the “Wilcoxon signed-rank test” since our data comes in the category of related samples. Multiple corrections need to be carried out to control for false discovery rate (FDR), which was performed using the method proposed in The control of the false discovery rate in multiple testing under dependency.

## 3. Results

From the above define experiments, 800200 p-values were obtained for eLORETA, LCMV, and BC-VARETA. Later these p-values were corrected for False Discovery Rate (FDR). For a conclusive and comprehensive analysis of corrected p-values, we plotted the histograms of these p-values for sources computed from all three inverse methods. shown in (Figure - 2, 3 & 4). It can be seen that all p-values are greater than 0.01 for histograms of p-values computed for eLORETA, LCMV, and BC-VARETA. For the rejection of the null hypothesis, we set the significance level at .01. We can see all p-values are greater than 0.01 with BC-VARETA having minimum p-value at 0.0414, eLORETA at 0.362, and LCMV at 0.2477. Looking at these p-values it can be safelty said that the null hypothesis (both modalities are the same) is accepted as all p-values are greater than the significance level.

**Figure 4:**
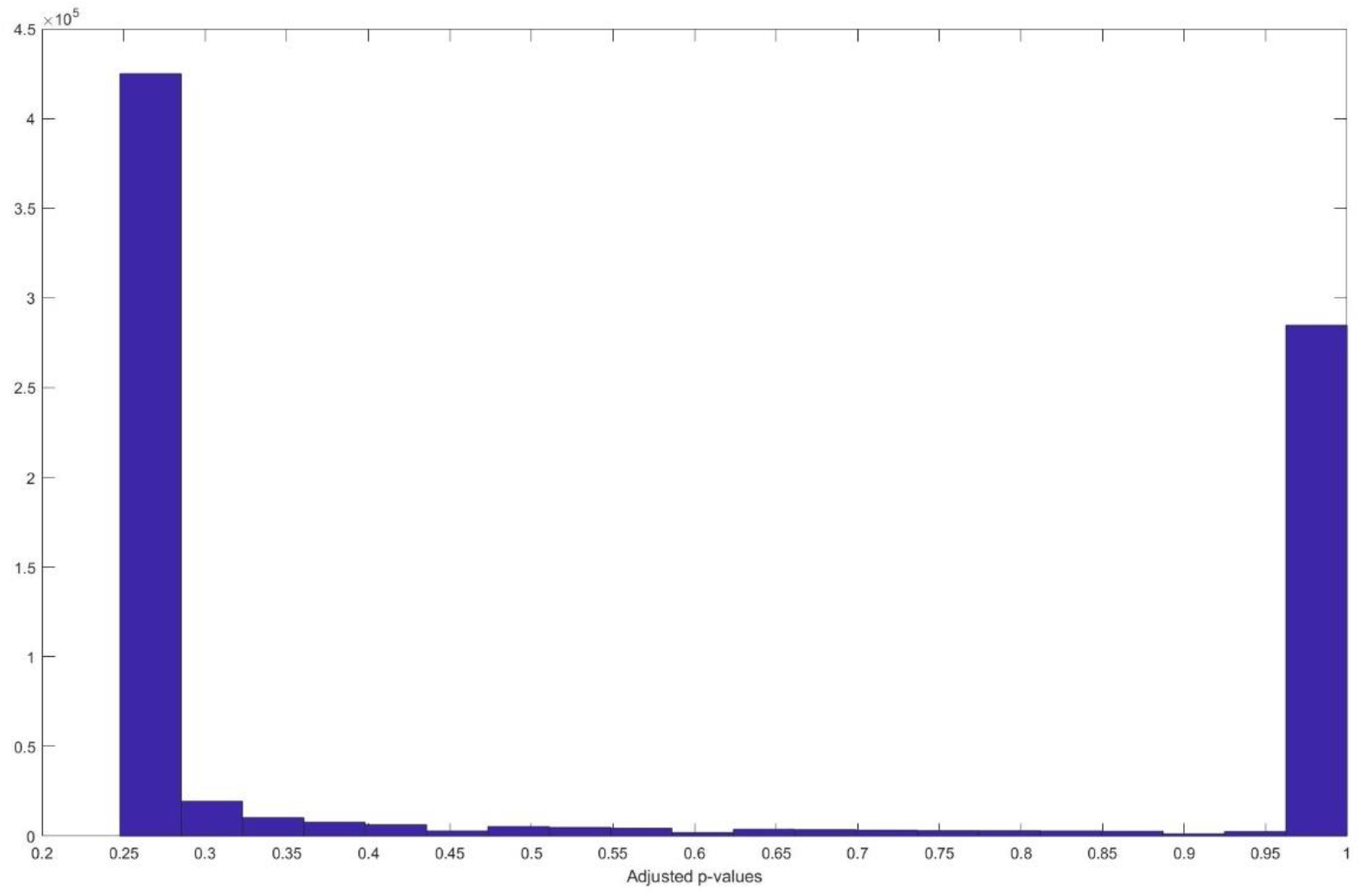
stogram of adjusted p-values for LCMV

**Figure 5:**
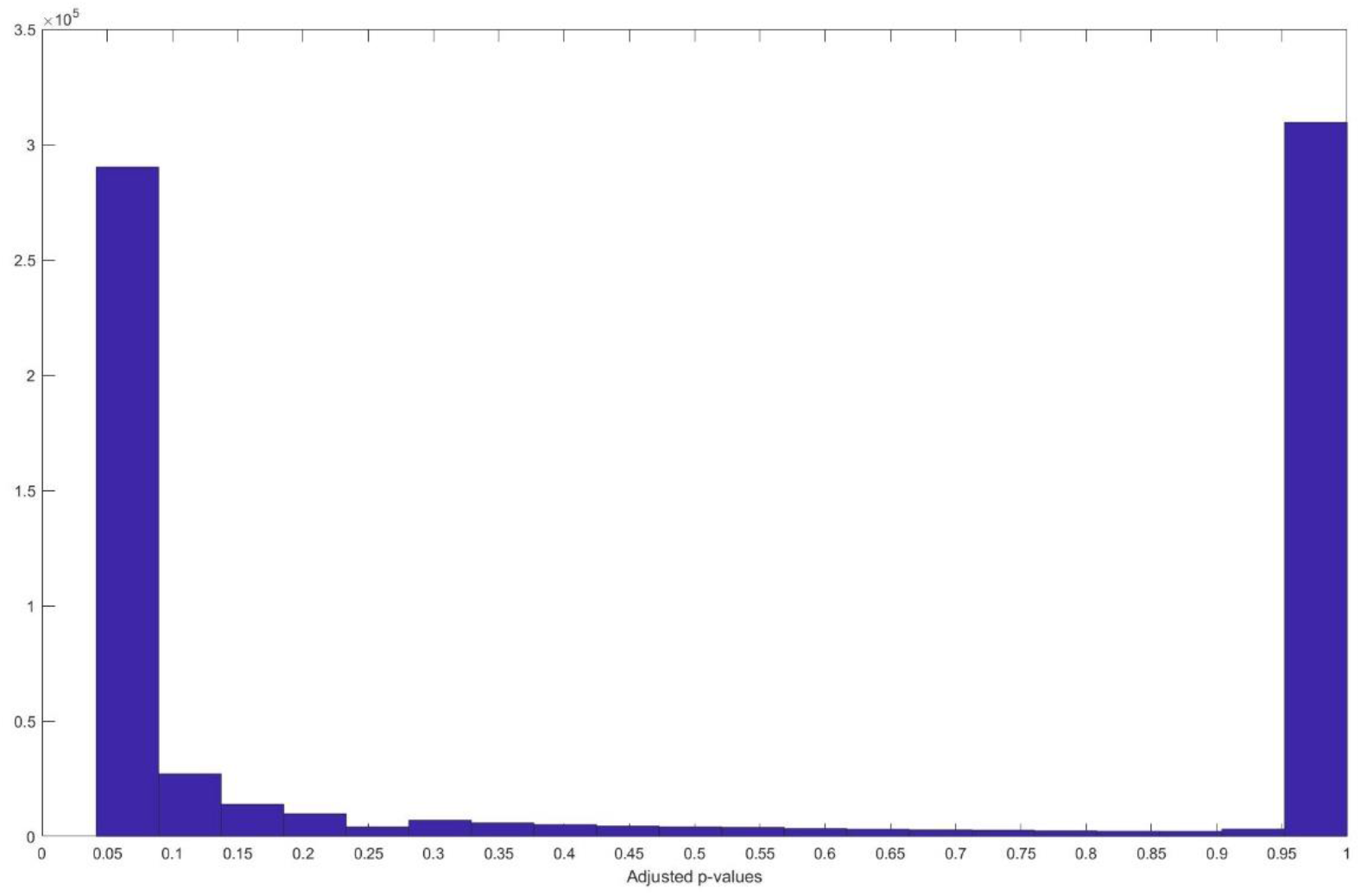
Histogram of adjusted p-values for BC-VARETA

## 4. Discussion

Since, EEG and MEG are capturing cortical activity in form of electric field and magnetic field generated by currents induced by same post-synaptic potentials (PSPs). So based on this underlying truth that both modalities are computing same activity, the sources computed from the inverse solutions should be same or not very much different. That is what we have analysed in this study by comparing EEG and MEG sources computed from two different datasets i.e. HCP-MEG and CHBM-EEG. We have analysed using permutation based rank test to see for any evidence that there are some differences in sources captured from both modalities. However, our results show that we don’t have any evidence to show any statistical differences between sources from two modalities. These finidings can have very innovative and optimistic implications in the field on neuroscience. Firstly, it counters the ideas that promotes to perform simultaneous EEG and MEG studies to have a better picture of cortical activity. Since there are not much differences in sources computed from both modalities, it can be safely said that studies performed based on either EEG or MEG will still give us detailed information of all underlying sources in cortex. Since EEG is easily portable and MEG is still under process of being transformed into a portable scanning setup. So studies in remote areas and also for underdeveloped or poor countries who cant afford expensive setup required for MEG can be carried out using EEG. Secondly, using sophisticated inverse solutions like we used in this studi, we can extract similar underlying cortical avtivity from EEG that is extracted from MEG.

In this study, we have compared anatomically the same sources, which means that EEG and MEG are capturing the same cortical activity i.e. scanning resting-state cortical activity in humans. Additionally, before making any comparisons we have corrected our source spectra for any possible sources of differences due to the engineering and physics of recording machines. Due to these reasons, the sources computed from inverse solutions should be the same or should have a close correlation between them. That is exactly what we have observed in our results once we compared the sources computed from BC-VARETA. The results have shown that when a permutation-based univariate rank test was carried out to make a comparison between EEG and MEG sources computed from the BC-VAERTA for 140 subjects showed no differences for any source or frequency points. There could have been some source of possible differences between the two modalities because of differences in frequency response, engineering, and design of amplifiers. To counter these differences we corrected sources spectra for amplifier differences before performing permutation tests. Another correction for global scale differences was required to be made for each subject to correct for any possible subject-based differences. A comparison would be fair once it is made after all the above-defined corrections. We have made all the corrections and found out no differences in both modalities. P-values computed from univariate permutation analysis were corrected for False Discovery Rate (FDR). The corrected p-values showed that our null hypothesis i.e. there are no differences between EEG and MEG is accepted with a confidence interval of 99%. To compare the findings of our study and make the analysis more concrete, we performed all the above-defined operations to EEG and MEG using state of the art inverse solutions i.e. eLORETA and LCMV. Our findings from these two methods were no different from those we found in BC-VARETA i.e. the rhythmic activity among sources of EEG and MEG are statistically similar.

Our research and results have stated that the source spectra of M/EEG are statistically the same, it contradict with the recent paper by Christian Benar at el. They provided some initial evidence that MEG and EEG differ in terms of background activity (C.G. Bénar, 2019). The results of our study lead to further analysis for the understanding of the physiological mechanism and activation state of M/EEG. However, the findings of our study give clear evidence that EEG, if performed an analyzed properly can give a detailed picture of cortical activity similar to MEG and vice versa. Results of this study imply that the conventions that are made from many studies that sources captured from EEG and MEG are different and some sources are better detected using one modality are wrong. EEG can be used to record sources if properly used.

## 5. Conclusion

To the best of our knowledge, this is the first study to perform a statistical comparison between the sources of EEG and MEG rhythms, with a large number of healthy subjects in resting state, using a novel inverse solution of BC-VARETA. This study also compares the results of BC-VARETA with state of the art inverse methods eLORETA and LCMV. Furthermore, to make an extensive and concrete comparison we have computed sources for complete spectra of EEG (0-50Hz) and similar frequency range for MEG. Before performing statistical comparison we have corrected source spectra of EEG and MEG for amplifier differences and global scale differences among individuals. For statistical comparison, a univariate permutation test was performed with additional FDR correction on each source and each frequency point variable of both modalities. Upon looking at the corrected p-values we have concluded that sources computed for both modalities using all three inverse solutions described are the same with a significance level of 0.01. This is a very important finding can help conducting further studies not only focused on simultaneous modalitiy studies. In fact cortial activity captured from either EEG or MEG can give detailed knowledge of underlying cortical activity given that condition that sophisticated and modern inverse solutions are applied. The possible future directions are: Comparing sources of simultaneous EEG and MEG rhythms and their connectivity and checking if we can get more stable statistical similarities.

## 6. Funding

The authors were funded from the University of Electronic Sciences and Technology grant Y03111023901014005 and from the National Science Foundation of China (NSFC) with the grant 61871105 and 81861128001.

## 7. Acknowledgments

The authors would like to thank for the support from the NSFC (China-Cuba-Canada) project (No. 81861128001) and the funds from National Nature and Science Foundation of China (NSFC) with funding No. 61871105, 61673090, and 81330032, and CNS Programme of UESTC (No. Y0301902610100201). The authors would like to acknowledge the Cuban Human Brain Mapping Project at the Cuban Neuroscience Center and the Ministry of Public Health of Cuba and WU-Minn Human Connectome Project that provide the EEG and MEG data. We also wish to thank all the members in the Joint China-Cuba Lab for Frontier in Translational Neurotechnology of the University of Electronic Science and Technology of China.

## 8. Data availability statement

The dataset (EEG) from Cuban Brain Mapping Project for this study is available on request to Denys Buedo Hidalgo or Iris Rodriguez.

The dataset (MEG) from the Human Connectome Project for this study can be accessed on request at https://www.humanconnectome.org/.

## Notes

### Competing Interest Statement

The authors have declared no competing interest.

### Summary of Updates

Abstract, Methodology, Results and Discussion updated

